# RTN4IP1 is essential for the final stages of mitochondrial complex I assembly and coenzyme Q biosynthesis

**DOI:** 10.1101/2024.09.04.610987

**Authors:** Monika Oláhová, Rachel M. Guerra, Jack J. Collier, Juliana Heidler, Kyle Thompson, Chelsea R. White, Paulina Castañeda-Tamez, Alfredo Cabrera-Orefice, Robert N. Lightowlers, Zofia M. A. Chrzanowska-Lightowlers, Ilka Wittig, David J. Pagliarini, Robert W. Taylor

## Abstract

A biochemical deficiency of mitochondrial complex I (CI) underlies ∼30% of cases of primary mitochondrial disease, yet the inventory of molecular machinery required for CI assembly remains incomplete. We previously characterised patients with isolated CI deficiency caused by segregating variants in *RTN4IP1,* encoding a mitochondrial NAD(P)H oxidoreductase. Here, we demonstrate that RTN4IP1 deficiency causes a CI assembly defect in both patient fibroblasts and knockout cells, and report that RTN4IP1 is a *bona fide* CI assembly factor. Complexome profiling revealed accumulation of unincorporated ND5-module and impaired N-module production. RTN4IP1 patient fibroblasts also exhibited defective coenzyme Q biosynthesis, substantiating this emerging function of RTN4IP1. Thus, our data reveal RTN4IP1 plays an essential role in both the terminal stages of CI assembly and in coenzyme Q metabolism, and that pathogenic *RTN4IP1* variants impair both functions in patients with mitochondrial disease.

## Introduction

Mitochondria are functionally diverse organelles whose importance in humans is most strongly evidenced by their dysfunction in patients with primary mitochondrial disease ^1–3^. This group of disorders demonstrates remarkable clinical, genetic, and functional heterogeneity, and is caused by pathogenic variants in genes encoding mitochondrial components ^4^. The largest group of mitochondrial diseases are caused by variants affecting oxidative phosphorylation (OxPhos) ^5^, the process by which mitochondria synthesise ATP. OxPhos is driven by five complexes (CI-V) embedded within the inner mitochondrial membrane. CI-IV comprise the mitochondrial respiratory chain, which generates the proton motive force coupled to the electron transfer through a series of redox centres, terminating in oxygen. CV harnesses this electrochemical proton gradient to drive ATP production ^6,7^.

OxPhos complexes are composed of multiple subunits and cofactors. Several mitochondrial assembly factors and membrane insertases ensure their faithful formation ^7^. CI (NADH:ubiquinone oxidoreductase), the largest respiratory chain complex, is a ∼1 MDa multimeric assembly of 45 different subunits encoded by genes in either the nuclear or mitochondrial genomes and is the primary entry point for electrons, which are transferred from NADH along a series of 7 iron-sulphur (Fe-S) clusters to ubiquinone (CoQ) ^8^. Biogenesis of CI is a stepwise process involving modular assembly coordinated by at least 18 known assembly factors ^9–11^. In humans, CI assembly requires the formation of distinct intermediate pre-assemblies to form a functional enzyme. The Q- and N-modules form the hydrophilic matrix arm of the enzyme whilst ND1-, ND2-, ND4- and ND5-modules produce the hydrophobic membrane arm. The insertion of mitochondrial DNA encoded CI subunits (MT-ND1, MT-ND2, MT-ND3, MT-ND4, MT-ND4L, MT-ND5 and MT-ND6) into the membrane arm modules appears to be regulated by specific co-translational systems, involving the mitochondrial insertase OXA1L and the translational regulator of CI, MITRAC15 ^12–14^. CI assembly is initiated by the union of the proximal membrane arm sub-assemblies, ND1 and ND2, in the inner mitochondrial membrane, and joining of the Q-module. The distal membrane arm ND4- and ND5-module sub-complexes join with the proximal membrane arm sub-assemblies before the final stages of CI assembly can take place. Finally, the N-module is incorporated by merging with the Q-module, thus completing the peripheral arm of CI ^10,15,16^. Fully assembled CI can form respiratory supercomplexes with complexes III_2_ and IV, however, the physiological role for these supercomplex assemblies continues to be debated ^17–20^.

CI deficiencies account for approximately 30% of all primary mitochondrial disease cases ^21,22^. Genomic and multiomic technologies continue to help identify causal disease-associated genetic variants and delineate the mechanisms regulating CI biology ^9,11,15,23,24^. Additional molecular genetic investigations of patients with CI deficiency have validated the role of candidate genes in CI assembly, critically shaping our understanding of this process ^25–27^. Despite these synergetic approaches, the complete repertoire of proteins required for CI production remains undetermined. Given the predominance of CI deficiencies in causing mitochondrial disease phenotypes, and the reported roles of CI in cancer ^28–30^ and neurodegeneration including Parkinson’s disease ^29,31,32^, it is essential to further improve the mechanistic resolution of CI assembly pathways and CI biology.

We previously identified a series of patients, including one with isolated CI deficiency, presenting with variable neurological phenotypes ranging from isolated optic atrophy to severe early-onset encephalopathy due to pathogenic, bi-allelic, variants in the mitochondrial Reticulon-4-Interacting Protein 1 (RTN4IP1) gene ^33^. Here, we demonstrate that *RTN4IP1* encodes a late-stage CI assembly factor and show that its roles in both CI assembly and CoQ biosynthesis are linked to disease pathology.

## RESULTS

### *RTN4IP1* patient fibroblasts and *RTN4IP1* knockout cells exhibit mitochondrial CI respiratory deficiency

Following the first reported association of bi-allelic, pathogenic *RTN4IP1* gene variants with either isolated optic atrophy or a multisystem neurological disease presentation with optic nerve involvement ^34^, (*OMIM: 610502*), targeted and whole exome sequencing approaches have been used to identify further disease-causing variants and establish a genotype-phenotype correlation ^34,33,35^. As part of the *RTN4IP1*-patient cohort reported by Charif and colleagues (2018), we identified a female child harbouring bi-allelic pathogenic variants in *RTN4IP1* (*NM_032730.5*); a maternally-inherited c.500C>T, p.Ser167Phe and a *de novo* c.806 + 1G>A variant which impaired splicing of *RTN4IP1* transcripts (**Figure 1A** and ^33^). Also known as Optic Atrophy-10 (OPA10), at the time RTN4IP1 was assigned as a mitochondrial uncharacterised protein (MXP) ^23^ that was later shown experimentally to be essential for normal OxPhos function ^36^. We showed that these specific *RTN4IP1* variants led to complete loss of immunodetectable RTN4IP1 protein, resulting in a CI assembly defect in patient skeletal muscle (Family 11 in ^33^).

**Figure 1.**
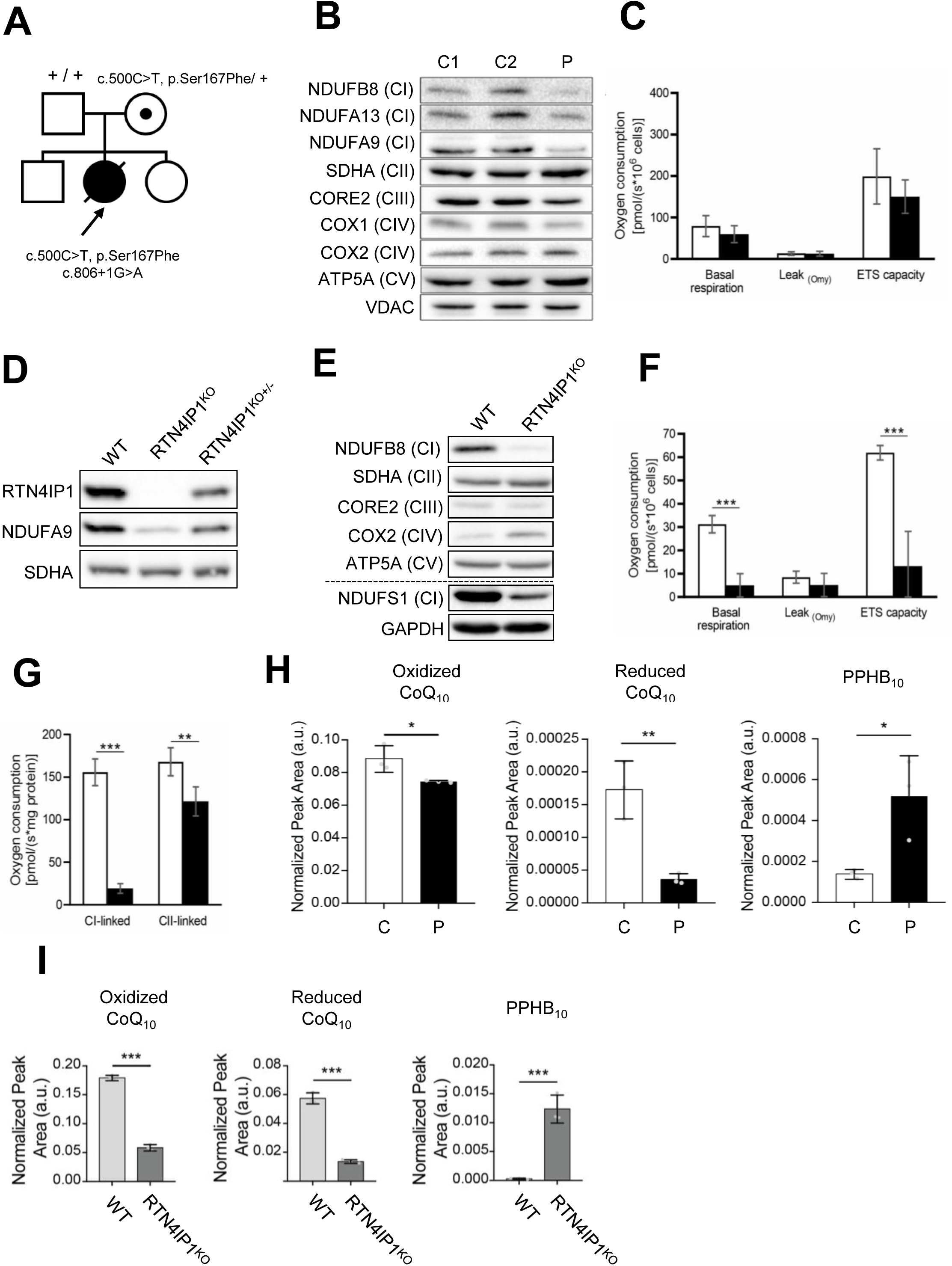
RTN4IP1 deficiency impairs CI protein levels, cellular respiration and CoQ biosynthesis. (**A**) Family pedigree showing pathogenic *RTN4IP1* variants. (**B**) Western blot analysis of protein extracts from control (C1, C2) and *RTN4IP1* patient fibroblasts (P) showing an isolated CI defect. VDAC and SDHA were used as loading controls (n=2). (**C**) High-resolution respirometry of patient-derived *RTN4IP1* fibroblasts (black bars) compared to control fibroblasts (white bars). ROX-corrected analysis of basal respiration, Oligomycin (Omy)-inhibited Leak respiration and maximum uncoupled respiration (ETS capacity) are shown. (n=3 mean ± SD). (**D**) Western blot analysis confirmed the generation of CRISPR-Cas9-edited heterozygous (+/-) and homozygous (-/-) RTN4IP1^KO^ U2OS lines and isogenic WT control. SDHA was used as loading control (n=2). (**E**) Western blot analysis of OxPhos complex subunits, showing and isolated CI defect in RTN4IP1^KO^ compared to WT. GAPDH and SDHA were used as loading controls (n=3). (**F-G**) Respirometry analysis as described in (**C**) in U2OS WT (white bars) and RTN4IP1^KO^ (black bars) cells showing a respiration defect caused by a severe reduction of CI-linked respiration. (n=4 mean ± SD). (**H-I**) Targeted lipidomic analysis on (**H**) *RTN4IP1* patient (P) and control (C) fibroblasts and (**I**) RTN4IP1^KO^ and WT U2OS cells, (n=3 mean ± SD). *p<0.05, **p<0.01, ***p<0.001, two-sided Student’s t-test.

To further interrogate the role of RTN4IP1 in mitochondrial metabolism, we characterised the OxPhos function in primary *RTN4IP1-*patient fibroblasts carrying c.500C>T; c.806 + 1G>A variants and in a CRISPR-Cas9 generated U2OS RTN4IP1 knockout cell model. SDS-PAGE and immunoblotting showed a marked steady-state decrease in the CI subunits NDUFB8, NDUFA13, and NDUFA9 in *RTN4IP1* patient fibroblasts. Subunits from OxPhos CII-CV were unaffected compared to controls (**Figure 1B**). Next, we investigated the respiratory capacity of *RTN4IP1-*patient fibroblasts, observing a mild decrease (not-significant) in basal respiration in patient-derived cells relative to control (**Figure 1C**). Oxygen consumption promoted by proton leakage upon CV inhibition was unaffected in *RTN4IP1* patient compared to control fibroblasts, signifying normal integrity of the inner mitochondrial membrane and coupling of respiration with ATP synthesis (**Figure 1C**). The electron transport system (ETS) capacity, representing uncoupled respiration at optimum carbonyl cyanide p-trifluoro methoxyphenylhydrazone (FCCP) concentration for maximum respiratory flux, was slightly decreased in the *RTN4IP1* patient, although this was not significant (**Figure 1C**).

To further investigate the molecular functions of RTN4IP1, we generated a tractable model cell system (U2OS) lacking RTN4IP1 to facilitate an extended characterisation of RTN4IP1 function. Using CRISPR-Cas9 genome editing, we created heterozygous RTN4IP1^KO+/-^ and homozygous RTN4IP1^KO-/-^ (hereinafter RTN4IP1^KO^) cell lines, as well as isogenic wild-type (WT) controls. RTN4IP1^KO+/-^ and RTN4IP1^KO^ cells showed decreased and completely abolished RTN4IP1 protein levels, respectively (**Figure 1D**). Similar to *RTN4IP1-*patient fibroblasts, immunoblotting revealed a marked decrease in the steady-state levels of assessed CI subunits NDUFA9, NDUFB8 and NDUFS1 in RTN4IP1^KO^ cells compared to WT (**Figures 1D and E**). Steady-state protein levels of all other assessed OxPhos components were largely unaffected (**Figure 1E**). Respirometry measurements in RTN4IP1^KO^ cells revealed a severe decrease in oxidative capacity (**Figure 1F**). While the proton leak-linked respiration rate, resulting from the addition of oligomycin, was not significantly different between WT and RTN4P1^KO^, ETS capacity was significantly diminished in the absence of RTN4IP1 compared to WT control (**Figure 1F**). To further analyse the underlying mechanism leading to decreased oxygen consumption in cells lacking RTN4IP1, CI- and CII-linked respiratory rates were measured, showing significant reductions of >88% and ∼28%, respectively. Our collective data point towards CI as the precise site responsible for this observed decrease in respiration (**Figure 1G**).

### Pathogenic *RTN4IP1* variants lead to an impairment in coenzyme Q biosynthesis

Previous characterisation of protein-protein interactions by affinity enrichment mass spectrometry (AE-MS) of 50 MXPs ^23^ captured an interaction between RTN4IP1 and COQ9, an auxiliary lipid-binding protein involved in the CoQ biosynthesis pathway ^37^. Furthermore, recently published work has characterized RTN4IP1 as a mitochondrial matrix oxidoreductase that supports coenzyme Q biosynthesis, specifically through regulation of the O-methyltransferase activity of COQ3 ^38^. We therefore performed a targeted lipidomic analysis of *RTN4IP1-*patient fibroblasts, which revealed a mild loss of oxidized coenzyme Q10 (CoQ_10_), and a more striking loss of reduced CoQ_10_ compared to control fibroblasts, which was accompanied by elevated polyprenyl-hydroxybenzoate 10 (PPHB_10_), an early CoQ biosynthesis pathway intermediate that typically accumulates when the CoQ pathway is defective (**Figure 1H**). Targeted lipidomics of both WT and RTN4IP1^KO^ cells revealed a significant decrease in both oxidized and reduced CoQ_10_ levels, accompanied by an increase in PPHB_10_ (**Figure 1I**). The marked loss of CoQ_10,_ together with the concomitant accumulation of an early biosynthetic intermediate, is consistent with RTN4IP1 regulating this pathway. To further probe the role of RTN4IP1 in CoQ biosynthesis, we generated stable overexpression of WT RTN4IP1 in the RTN4IP1^KO^ cell line (**Figure SI 1A**) and performed lipidomic analyses. Stable overexpression of RTN4IP1 partially rescued the CoQ deficiency in the RTN4IP1^KO^ cell line (**Figure SI 1B**), further underscoring the physiological role of RTN4IP1 in supporting efficient CoQ_10_ biosynthesis.

### RTN4IP1 deletion selectively impairs CI biogenesis

A genome-wide CRISPR-based knockout screen previously identified RTN4IP1 as a candidate OxPhos disease gene with an unassigned function ^36^. Given the severe CI-related defects documented in the RTN4IP1^KO^ line (**Figure 1D-E**), in addition to those described in *RTN4IP1*-patient tissues (**Figure 1B**) ^33^, we performed proteomic analyses of WT and RTN4IP1^KO^ cells to further define the role of RTN4IP1 in CI biology. Absence of RTN4IP1 caused a decrease in the abundance of CI subunits, whilst the levels of CII, CIII, CIV and CV components were largely unaffected (**Figures 2A-B**). Consistent with this, gene ontology overrepresentation analysis of the full proteomic profile of RTN4IP1^KO^ cells revealed enrichment of factors involved in NADH dehydrogenase complex assembly (GO: 0010257) among the most downregulated proteins (**Figure SI 2A**). Stable overexpression of RTN4IP1 (RTN4IP1^FLAG^) restored the levels of most downregulated CI subunits in the RTN4IP1^KO^ cell line closer to WT levels (**Figures SI 2B-C**), demonstrating that defective RTN4IP1 causes CI-deficiency.

**Figure 2.**
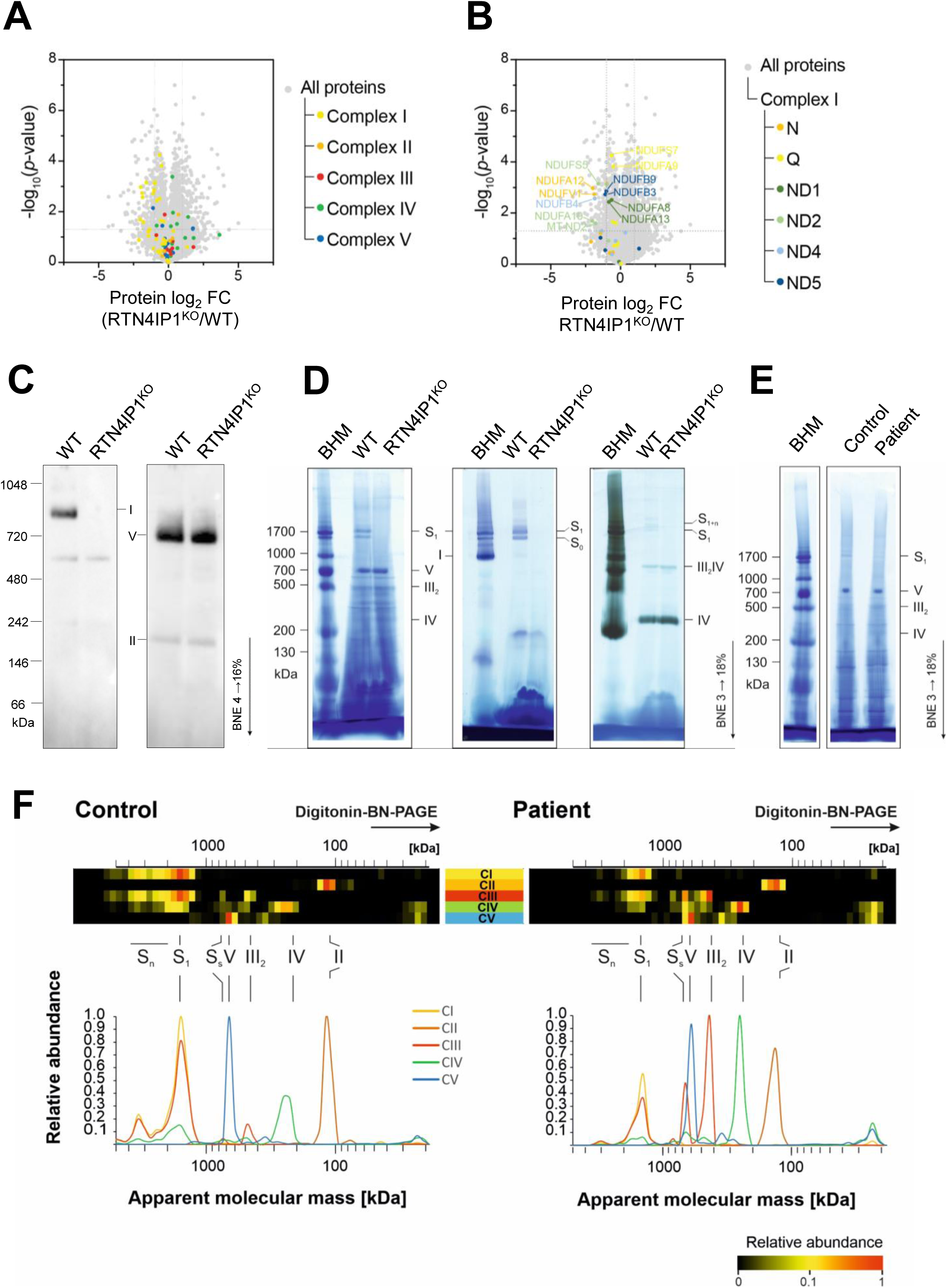
RTN4IP1 deficiency selectively impairs CI biogenesis. (**A-B**) Volcano plot of (**A**) CI-V protein abundances in U2OS RTN4IP1^KO^(*+GFP*) cells relative to WT(*+GFP*) and in (**B**) the decrease in distinct CI subunits following loss of RTN4IP1 are shown. In (**B**) CI proteins are highlighted, and colour coordinated to match individual CI modules. Data shown as mean (n=3), two-sided Student’s t-test. (**C**) BNE and immunoblotting analysis of DDM solubilised mitochondrial membranes from WT and RTN4IP1^KO^ U2OS lines using antibodies against complexes I (NDUFV1), II (SDHA, loading control) and V (ATP5A), (n=3). (**D-E**) In (**D**) deletion of RTN4IP1 results in absence of supercomplex and undetectable NADH:NTB reductase activity. Digitonin solubilisation of mitochondrial membranes from U2OS WT and RTN4IP1^KO^ cells separated on native gradient gels were stained with Coomassie (left), NADH:NTB reductase activity stain (middle) and CIV specific heme stain (right). Representative gels for NADH:NTB and heme stain are shown (n=3). In (**E**) *RTN4IP1* variants cause reduction of CI-containing supercomplexes. Enriched mitochondrial membranes from control and the *RTN4IP1* patient fibroblasts were solubilised with digitonin and separated on a native gradient gel. Protein complexes were stained with Coomassie, (n=3). (**F**) Summary of OxPhos complexes from complexome profiling of control and *RTN4IP1* patient fibroblasts shown as 2-D OxPhos complexes profiles. Subunits of each individual OxPhos complex were summed up and normalized to maximum between gel lanes of control and patient samples and illustrated in a heatmap. Assignment of complexes in (**D-F**): complex I (I); complex II (II); complex III dimer (III_2_); complex IV (IV); complex V (V); supercomplex containing CI, III_2_ and 1 copy of CIV (S_1_); S_1_ and extra copies of CIV (S_1+n_); higher order supercomlexes (S_n_). BHM - bovine heart mitochondria solubilized with digitonin served as native mass ladder.

Given the observed decrease in CI subunits in RTN4IP1-deficient cells, we next investigated how loss of RTN4IP1 affected the assembly of OxPhos complexes by analysing N-dodecyl β-D-maltoside (DDM)-solubilised mitochondrial membrane extracts from WT and RTN4IP1^KO^ using blue native electrophoresis (BNE) and immunoblotting analysis. Applying antibodies against the N-module subunit of CI, NDUFV1, we were unable to detect fully assembled CI in RTN4IP1^KO^ compared to WT, indicating a key role for RTN4IP1 in CI biogenesis (**Figure 2C**), which is in line with the decrease in steady state CI subunits in the RTN4IP1^KO^ proteomics data (**Figures 2A-B**).

Next, we solubilised mitochondrial membranes from WT and RTN4IP1^KO^ cells with digitonin. BNE analysis of Coomassie-stained native gels showed a clear absence of supercomplex S_1_ (I-III_2_-IV) in RTN4IP1^KO^ compared to WT (**Figure 2D**, left) and undetectable S_1_ when stained for NADH dehydrogenase activity (**Figure 2D**, middle). Moreover, heme staining showed that the pattern of CIV was not affected by loss of RTN4IP1 (**Figure 2D**, right). As expected, digitonin-solubilised mitochondrial membranes from *RTN4IP1*-derived patient fibroblasts also showed a clear reduction of the CI-containing supercomplex S_1_ compared to control following BNE (**Figure 2E**).

To define the RTN4IP1-associated OxPhos defect with higher resolution, we performed complexome profiling on the patient-derived cells and RTN4IP1^KO^ cells ^39^. Mitochondria were solubilised with digitonin, followed by protein separation by BNE, fractionation and analyses by quantitative MS as previously reported ^40^. Complexome profiling data were analysed to correlate differences in the abundance and arrangement of multiprotein complexes — from high (left) to low (right) molecular mass — in the form of a “heat map”, focussing specifically on the inventory of subunits and assembly factors of mitochondrial OxPhos components in *RTN4IP1* patient fibroblasts and RTN4IP1^KO^ cells and corresponding controls (**Figures 2F**, **3**, **4A, SI 3 and SI 4**). Consistent with the BNE gel data, supercomplex assembly (predominantly S_1_; I-III_2_- IV) was attenuated in *RTN4IP1-*patient cells (**Figures 2F and SI 3**), whilst a complete loss of fully assembled supercomplexes was noted in RTN4IP1^KO^ cells (**Figure 3**). Focussing on the RTN4IP1^KO^ cells, which demonstrated a more dramatic loss of CI-containing supercomplexes and CI-subcomplexes compared to *RTN4IP1* patient-derived fibroblasts, we observed diminished amounts of supercomplex S_1_ and the presence of a large intermediate of CI (L_int_) (**Figure 3B**, yellow box). This resulted in a concomitant increase of free CIII and CIV (**Figure 3B**, red box and green box, respectively). Similar increases in CIII and CIV were also observed in *RTN4IP1*-patient fibroblasts (**Figure SI 3,** red box and green box, respectively), whilst assembly of CII and CV were largely unaffected (**Figure SI 3)**. Collectively, these data support RTN4IP1 having an essential and specific role in CI assembly.

**Figure 3.**
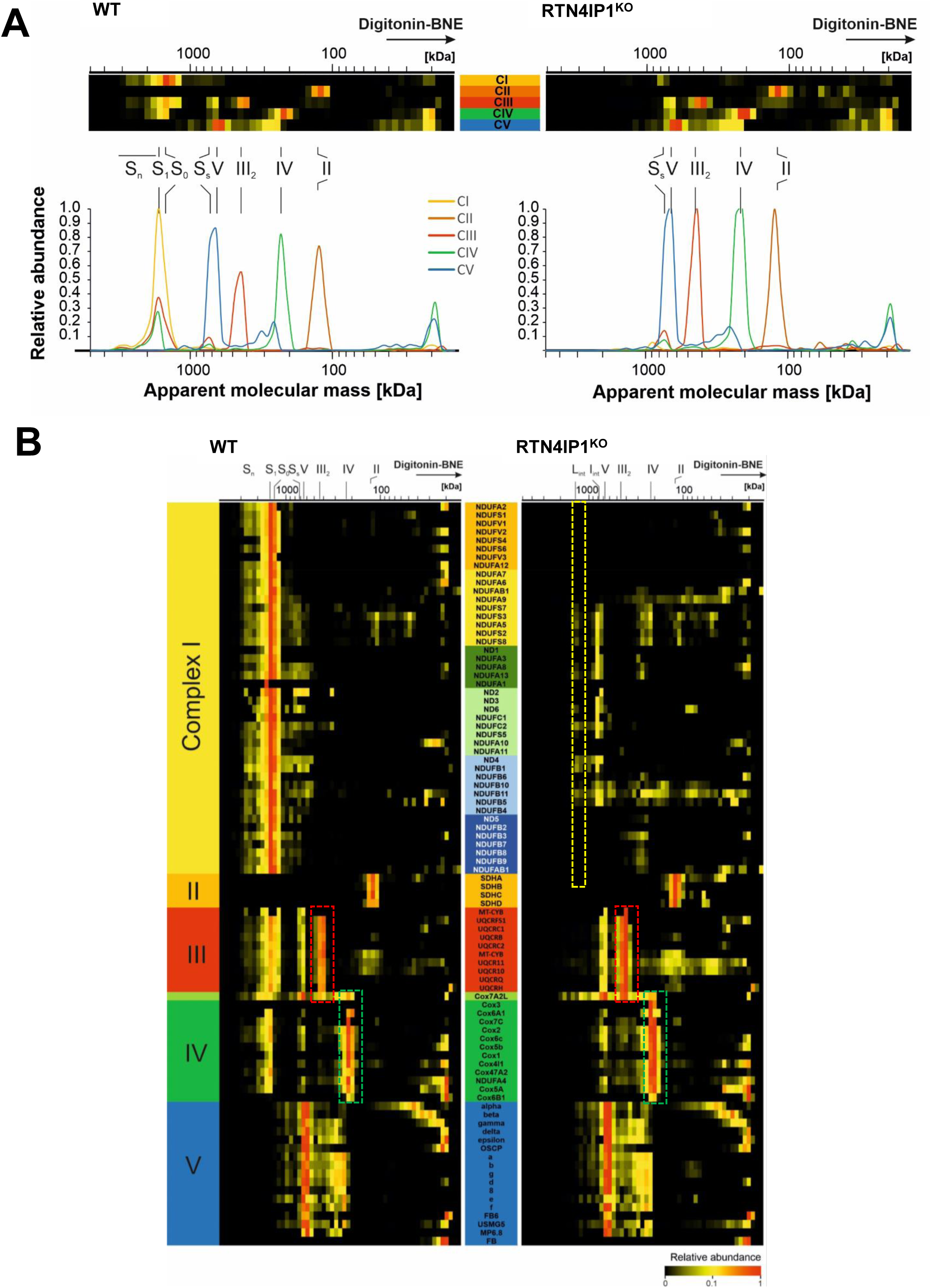
Complexome profiling analysis of OxPhos complexes from RTN4IP1^KO^ and WT U2OS cells. (**A**) 2-D OxPhos complex profiles from complexome data from RTN4IP1^KO^ corresponding to WT cells. Complexome profiling data of OxPhos complexes I-V are presented as 2-D plots. (**B**) OxPhos complexes of RTN4IP1^KO^ and WT cells analysed by complexome profiling. Complexome profiling data were presented as heat map, corresponding to subunits of individual OxPhos complexes I-V. In (**A-B**) Mitochondrial complexes were solubilized with digitonin, separated by BNE, and gel slices were analysed by qualitative MS. Assignment of complexes in (**A-B**): complex I (I); complex II (II); complex III dimer (III_2_); complex IV (IV); complex V (V); higher order supercomplexes (S_n_); supercomplex containing CI, III_2_ and 1 copy of CIV (S_1_); supercomplex containing CI and a dimer of CIII (S_0_); small supercomplex S_0_ of CIII_2_ and CIV (S_s_) and large intermediate CI (L_int_) and intermediate of CI including modules Q, ND1m, ND2m, ND4m and assembly factors (I_int_). The relative abundance of each protein was represented from low to high according to the colour scale illustrated on the bottom right.

**Figure 4.**
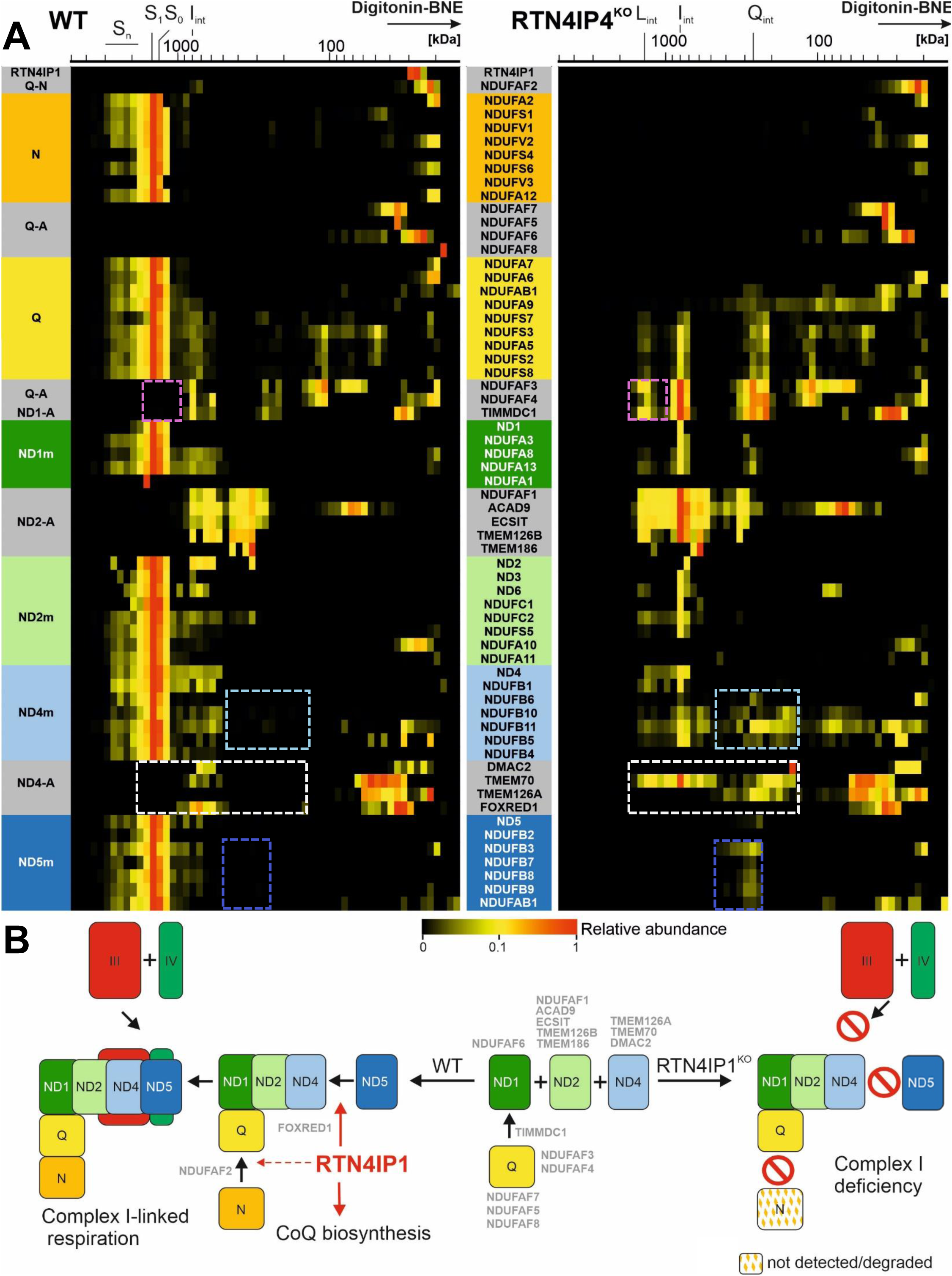
RTN4IP1 deletion impairs ND5 module attachment and N-module stability in the late stages of CI assembly. (**A**) Complexome profiling data of CI and identified CI assembly factors were sorted to their assembly modules and presented as heatmap. Assignment of complexes: higher order supercomplexes (S_n_); supercomplex S_1_ of CI, CIII_2_ and CIV (S_1_); supercomplex containing CI and a dimer of CIII (S_0_); large intermediate CI (L_int_); intermediate of CI including modules Q, ND1m, ND2m, ND4m and assembly factors (I_int_), intermediate of Q-module containing assembly factors (Q_int_). The relative abundance of each protein was represented from low to high according to the colour scale illustrated on the bottom. (**B**) Proposed role of RTN4IP1 in CI assembly. Modules Q (yellow), ND1 (dark green), ND2 (light green), ND4 (light blue) and ND5 (dark blue) are pre-built using assembly factors (grey). Individual modules assemble under the control of additional assembly factors shown in the centre (grey). Under WT conditions (left), the membrane arm is complete, allowing the N-module (orange) to attach to form a fully active CI enzyme. Complexes III (red) and IV (green) are also added to fulfil the function of CI-dependent respiration as a supercomplex. The conditions under loss of RTN4IP1 i.e. in the RTN4IP1^KO^ cells are shown on the right.

### RTN4IP1 is a late-stage CI assembly factor

To position RTN4IP1 within the stepwise CI biogenesis process, which involves the tightly coordinated assembly of intermediate modules (ND1, ND2, ND4, ND5, Q, and N) and cofactors (e.g., FMN, Fe-S clusters), we further interrogated the complexome profile of CI. Analysis of the complexome profile of both *RTN4IP1* patient fibroblasts and RTN4IP1^KO^ cells (**Figures 4A and SI 4**) demonstrated that while the initial Q-module assembly appeared to be normal, its subsequent sub-assemblies containing membrane subunits clearly accumulated in the absence of RTN4IP1. We also observed the presence of the Q-module assembly factors NDUFAF3, NDUFAF4 and TIMMDC1, which did not dissociate from the large intermediates (**Figure 4A**, purple box). ND1-module assembly was normal in RTN4IP1^KO^ cells and accumulated together with the Q- and mitochondrial Complex I assembly (MCIA) complex, which is important for the stability of mtDNA-encoded ND2 ^41^. Similarly, ND2-module assembly appeared normal as the ND2-module was found to co-migrate with Q- and ND1- modules and the MCIA complex (NDUFAF1, ACAD9, ECSIT, TMEM126B, TMEM186) (**Figure 4A**). Although the ND4-module assembled and joined with the Q-, ND1-, ND2-modules and the MCIA complex, it accumulated in sub-assemblies in RTN4IP1-deficient cells, involving at least five ND4-module subunits (NDUFB5, NDUFB6, NDUFB10 and NDUFB11) (**Figure 4A**, light blue box) and the assembly factors TMEM70 and TMEM126A (**Figure 4A**, white box). It is noteworthy that WT cells also exhibited the presence of a CI intermediate (I_int_), which encompasses the modules Q/ND1/ND2/ND4 in conjunction with assembly factors. We observed that FOXRED1, which was recently identified together with other assembly factors (TMEM70, DMAC1, DMAC2, TMEM126A) in ND4 intermediates ^11,15,42^, binds to I_int_ in WT, but appears to be absent in the intermediate I_int_ and L_int_ of RTN4IP1-deficient cells (**Figure 4A**, white box), suggesting that the absence of RTN4IP1 may disrupt the function of FOXRED1 in late CI membrane arm assembly. The most dramatic changes observed in the absence of RTN4IP1 were related to the assembly of ND5-module, which assembled only as a single intermediate and was not able to join to the ND4-module and the large Q/ND1/ND2-intermediate (**Figure 4A**, dark blue box). We did not observe an accumulation of N-module subunits at higher molecular masses in the RTN4IP1^KO^ cells, leading us to hypothesise that stalling of the CI assembly pathway at this stage could result in increased turnover of the N-module to avoid deleterious redox effects ^43^ (**Figure 4A**).

### RTN4IP1 deficiency impairs CI assembly in mitochondrial disease

Similar to RTN4IP1^KO^ cells, we found increased Q-module sub-assemblies retaining assembly factors and accumulating in *RTN4IP1*-patient fibroblasts when compared to controls. In addition, a comparable large assembly intermediate I_int_ including Q/ND1/ND2/ND4-modules — just as described in the RTN4IP1^KO^ cells — was present in the *RTN4IP1* patient cells **(Figure SI 4**). In contrast to RTN4IP1^KO^ cells, we observed that the ND5-module and the final N-module do assemble to complete CI and integrate into supercomplexes. However, we believe this could be the limiting step in this process, as all residual sub-assemblies (I_int_) appear to accumulate with their respective assembly factors (**Figure SI 4**). Finally, RTN4IP1 was only identified as an individual protein that did not migrate with any of the CI-pre-assemblies when digitonin treated mitochondrial membranes were separated by BNE and analysed by complexome profiling (**Figures 4A and SI 4**). Collectively, our data converge to show that RTN4IP1 is a CI assembly factor required for the terminal stages of CI assembly (**Figure 4B**).

## DISCUSSION

Here, we report a critical role for RTN4IP1 in the late-stage assembly of mitochondrial CI and confirm its role in regulating CoQ_10_ biosynthesis in humans. Our interest in RTN4IP1 started with the investigation and subsequent molecular diagnosis of a patient with bi-allelic *RTN4IP1* variants who presented with a severe and isolated CI biochemical deficiency in muscle and fibroblasts ^33^. A closer inspection of the clinical phenotypes associated with primary CoQ_10_ deficiency and patients with *RTN4IP1*-related mitochondrial disease — and other CI-driven pathologies — reveal significant overlap.

Recessively inherited, pathogenic variants in the *RTN4IP1* gene were first identified in several families with isolated optic atrophy ^34^, leading to the characterisation of larger patient cohorts with variable neurological phenotypes, thus expanding the clinical phenotype of RTN4IP1-related mitochondrial disease ^33^. Optic atrophy and optic neuropathy are commonly associated with other mitochondrial disease pathologies implicating CI deficiency, including Leber hereditary optic neuropathy ^44^ as well as recessively-inherited variants in structural components of CI such as NDUFS6 ^45^, the CI assembly factor TMEM126A ^42^ and DNAJC30 which mediates CI repair ^46^. It is interesting to note that some patients with inherited pathogenic variants in genes implicated in primary CoQ_10_ deficiency manifest with optic atrophy as a feature of complex neurological phenotypes, including mutation of COQ1/PDSS1 ^47,48^, COQ2 ^49^ and COQ6 ^50^. We demonstrated that, in addition to CI deficiency, the CoQ biosynthesis pathway is also impaired in the *RTN4IP1* patient fibroblasts (**Figure 1G**). Recent studies, using immortalized C2C12 *RTN4IP1-KO* mouse myoblasts and a muscle-specific *dRTN4IP1-KD* fruit fly model, have shown that CoQ_2_ supplementation was unable to fully rescue the respiratory defect caused by RTN4IP1 deficiency ^38^, suggesting that the function of RTN4IP1 in CI assembly may be distinct from its role in CoQ_10_ biosynthesis, at least in mice and flies.

The biogenesis of CI is a complex and precisely orchestrated process, involving the assembly of pre-produced modules with the aid of specific CI assembly factors. We now identify RTN4IP1 as a novel CI assembly factor that completes the late stages of mitochondrial CI assembly. Tracking the late-acting CI assembly factors of the distal membrane part ND4/ND5-module bound to intermediates (**Figure 4A**, white box), we found that TMEM70 ^51^ and TMEM126A ^42^ are still involved in this process in the absence of RTN4IP1. However, DMAC2 ^11^ and FOXRED1 ^52^ have dissociated, as they are not found in high molecular mass intermediates and were only identified at the low molecular mass. Although the role of FOXRED1 in the late assembly of the membrane portion is established through the study of *FOXRED1* deficient cells, its specific molecular function as a putative FAD-dependent oxidoreductase is not completely elucidated ^52,53^. Based on our data we hypothesise that the mitochondrial matrix NAD(P)H oxidoreductase RTN4IP1 has a function at this point of CI assembly, as the termination products are found after the assembly of the ND4-module. Despite an exhaustive examination of the complexome data, deposited in the Proteomics IDEntifications (PRIDE) Archive database ^54^, RTN4IP1 was not identified in the high molecular mass region of the late assembly intermediates. This strongly suggests that RTN4IP1 may bind to the complex in a loose and transient manner, acting as a soluble matrix protein to fulfil its function as a late CI assembly factor. It remains unclear whether RTN4IP1 directly interacts with the ND4- or ND5-modules, or whether it functions as an oxidoreductase with FOXRED1, allowing the joining of ND4- and ND5-modules and signalling the readiness for N-module docking.

The ND4- and ND5-modules are important for the docking of other respiratory chain complexes. Inherited human CI deficiency associated with impaired assembly of very late CI subunits, such as NDUFA6 (N-module), showed that assemblies with complexes III and IV are still present in the patient cells, despite the lack of N-module^55^. Interestingly, the connection to complexes III and IV does not occur with the incomplete CI membrane arm in RTN4IP1^KO^ cells. This suggests that the necessary interfaces for supercomplex formation are indeed in the membrane arm, with NDUFA11 possibly playing an essential role at the interface with CIII ^56^. Although the CI assembly defect in the *RTN4IP1* patient fibroblasts is not as dramatic as in RTN4IP1^KO^ cells, there is a marked reduction in the preformed ND5-module and CI-containing supercomplex assembly when compared to controls (**Figures SI 3 and 4**).

RTN4IP1 is the second example, along with PYURF, of a protein working to connect the interrelated pathways of CI assembly and CoQ biosynthesis and it’s interesting that the only reported case of pathogenic human *PYURF* variants presented neonatally with profound metabolic acidosis and multisystem mitochondrial disease which included optic atrophy ^24^. The need for concerted regulation of both CI and CoQ biosynthesis could arise from different sources. First, it is plausible that such proteins are required at the interface of these two essential pathways to carefully tune the ratio of CoQ to CI for the specific redox requirements of OxPhos. Second, CoQ or intermediates in the biosynthetic pathway could be required as participants in the assembly process of CI. This hypothesis seems less well supported due to different reports of the activity and stability of CI in various mouse models of CoQ deficiency ^57,58^. Discriminating between these two possibilities would benefit from a rigorous assessment of assembled CI abundance in diverse models of CoQ deficiency. Although the mechanism by which PYURF coordinates CoQ biosynthesis and CI assembly is unclear, it is interesting to note that PYURF is reported to interact with COQ3, COQ5, and NDUFAF5, all methyltransferases in their respective pathways ^24^. Similarly, recent proximity interaction studies identify COQ3, COQ5, PYURF and NDUFAF7 as interacting partners of RTN4IP1 ^38^. Curiously, NDUFAF7, a CI assembly factor, has been characterized as a protein methyltransferase that is required to methylate NDUFS2 in order to facilitate early CI assembly ^59^. It is therefore tempting to speculate that, like PYURF, RTN4IP1 enables the independent co-regulation of CoQ biosynthesis and CI assembly by facilitating critical methyltransferase reactions in both pathways.

In summary, we have shown that the NAD(P)H oxidoreductase RTN4IP1 is essential for normal CoQ production in humans, and that it functions as a late-stage assembly factor, crucial for the stability of the ND5-module and its docking to the ND4-module, which is a prerequisite for the final steps of CI assembly.

## Materials and Methods

### Ethical approval

Informed consent for diagnostic and research studies was obtained in accordance with the Declaration of Helsinki protocols and approved by the Newcastle and North Tyneside Local Research Ethics Committee (REC ref 2002/205).

### Mammalian cell culture

Primary age-matched control and *RTN4IP1* patient fibroblasts and CRISPR-Cas9-edited U2OS RTN4IP1^KO+/-^, RTN4IP1^KO-/-^ and wild type (WT) cell lines were grown in high glucose Dulbecco’s modified Eagle’s medium (DMEM, Gibco™ 31966047) supplemented with 10% fetal bovine serum (Gibco™ 16140071), 1% penicillin-streptomycin (Gibco™ 15140122), 1x non-essential amino acids (Gibco™ 11140068), and 50 µg/mL uridine (Sigma U3003) at 37°C and 5% CO_2_.

### Generation of RTN4IP1^KO^ cell models

CRISPR-Cas9 genome editing was used to generate RTN4IP1^KO^ and isogenic control cell models. Online CRISPR design tools were used to identify a 20bp sequence single guide RNA (sgRNA) complementary to the target sequence in the *RTN4IP1* gene (GGATTAGTACTACCTCTCCT). The *RTN4IP1* sgRNA was cloned into CRISPR-Cas9 expression vector px459 (Addgene). Wild type U2OS cells were sub-cultured two days before nucleofection to achieve 80% confluency. Cells (1×10^6^) were nucleofected with the px459 expressing the 20bp *RTN4IP1* sgRNA according to manufacturer’s instructions (Lonza). Subsequently, cells expressing the px459 + *RTN4IP1* sgRNA plasmid were enriched by antibiotic selection for 48 hours, and individual single clones were expanded for two weeks. Genomic DNA was isolated from expanded clones by incubating cell pellets in the presence of DirectPCR Lysis Reagent (Mouse Tail) (Viagen) and Proteinase K (Sigma) at 56°C for 16 hours followed by 95°C for 15 min. A target region containing the edited RTN4IP1 site has been PCR amplified (primers upon request) and Sanger sequenced. Isogenic wild type control, heterozygous (c.112C>T / +) and homozygous (c.111_127del/c.111_116delinsTCAA) RTN4IP1^KO^ cell lines were selected and used for subsequent experimental analysis.

### Western blot analysis

Cell lysis was performed on frozen cell pellets using buffer containing 50 mM Tris-HCl pH 7.5, 130 mM NaCl, 2 mM MgCl_2_, 1 mM Phenylmethylsulphonyl fluoride (PMSF), 1% Nonidet™ P-40 (v/v) and 1x EDTA free protease inhibitor cocktail on ice for 20 min. Lysed samples were centrifuged at 1000 g for 5 min at 4°C. Soluble protein fractions were retained and analysed by Western blotting. Equal amounts of protein extracts (minimum 40µg) were electrophoretically separated on a 12% SDS polyacrylamide gel, followed by a wet transfer onto Immobilon-P Polyvinylidene Fluoride (PVDF) membrane (Merck Millipore) using the Mini Trans-Blot Cell system (BioRad).

### Immunoblotting

Primary and species appropriate HRP-conjugated secondary antibodies (Dako) were used; RTN4IP1 (Sigma SAB1408126), NDUFB8 (Abcam ab110242), SDHA (Abcam ab14715), UQCRC2 (Abcam ab14745), COXI (Abcam ab14705), ATP5A (Abcam ab14748), Total OxPhos Human WB Antibody Cocktail (ab110411), NDUFA9 (Molecular Probes, A21344), NDUFS1 (Proteintech 12444-1-AP) and GAPDH (Proteintech 60004-1-Ig). SuperSignal™ West Pico PLUS Chemiluminescent Substrate (Thermo Scientific) and the ChemiDoc (BioRad) system supported by the Image Lab software (BioRad) were used for visualization.

### Oxygen consumption

Mitochondrial basal respiration of intact cells was measured using high-resolution respirometry (Oxygraph-2k, Oroboros Instruments, Innsbruck, Austria) with DatLab software 6.1.0.7 (Oroboros Instruments, Innsbruck, Austria). Measurements were performed at 37°C in full growth medium (DMEM, 15% FCS, pyruvate, uridine, non-essential amino acids, penicillin/streptomycin). Basal respiration (Basal) was measured for 20 min followed by titration of oligomycin (2.5 µM final concentration f.c.) to measure oligomyin-inhibited LEAK respiration. Subsequently, uncoupler FCCP (Sigma Aldrich, Munich, Germany) was titrated stepwise (0.5 µM per step, f.c.) until maximal uncoupled respiration (ETS) was reached. Residual oxygen consumption (ROX) was determined after sequential inhibition of CIII with antimycin A (2.5 µM f.c.) and CIV with KCN (2 mM f.c.). To analyse the effect of RTN4IP1 deletion on CI– and CII-linked respiration isolated mitochondria were used. CI-and CII linked mitochondrial respiration was measured using high-resolution respirometry in MiR05-respiration media (110 mM sucrose, 60 mM potassium lactobionate, 0.5 mM EGTA, 3 mM MgCl_2_, 20 mM taurine, 10 mM KH_2_PO_4_, 20 mM Hepes, 2 mg/ml bovine serum albumin, pH 7.1) at 37 °C. Mitochondria were added to the chamber followed by the addition of the following substrates and inhibitors (final concentration): 2 mM malate,

2.5 mM pyruvate, 10 mM glutamate, 2.5 mM ADP, 0.5 µM rotenone, 10 mM succinate and 2.5 µM antimycin A. Absolute respiration rates were corrected for ROX and normalized to the total number of cells per chamber, to the protein content or to citrate synthase activity.

### Generation of Stable Cell Lines

Constructs of interest were cloned into pLVX-AcGFP1-N1 (Takara 632154) lentiviral vector under the CMV promoter using Gibson assembly. See Supplementary Table 1 for the constructs and oligonucleotides used in this study. Lentiviral particles were produced using the Lenti-X™ 293T system (Takara 632180). Viral media was collected 48 hours post-transfection and processed with a Lenti-X concentrator (Takara 631231) then centrifuged (4 hr, 600 x g, 4°C) before aliquoting, flash freezing, and storing at −80 °C. To transduce cells, U2OS RTN4IP1^KO^ cells were seeded in six-well plates, in the absence of penicillin-streptomycin, and incubated overnight. Media was supplemented with 0.5 μg/mL polybrene (Sigma TR-1003) and 20 μl of lentivirus was added to each well. Cells were incubated for 48 hours. The media was then exchanged for DMEM with 10% FBS and 0.5 μg /mL puromycin (Thermo A1113803) for 5 days. Cells were then passaged and expanded as normal and frozen down using freezing media (90% DMEM, 10% DMSO).

### Targeted CoQ Measurement by LC-MS/MS

Determination of the CoQ content and redox state in mammalian cell culture was performed as previously described with modifications ^60^. In brief, frozen cell pellets were resuspended in 100 μL of PBS and 5 μL was retained for a BCA assay to normalize lipidomic measurements to protein content. The remaining cell resuspension was added to ice cold extraction solution (600 μL acidified methanol [0.1% HCl final], 250 μL hexane, with 0.1 μM CoQ_8_ internal standard [Avanti Polar Lipids]). Samples were vortexed and centrifuged (5 min, 17000 x g, 4°C) and the top hexane layer was retained. Extraction was repeated twice before the hexane layers were combined and dried under argon gas at room temperature. Extracted dried lipids were resuspended in methanol containing 2 mM ammonium formate and overlaid with argon.

LC-MS analysis was performed using a Thermo Vanquish Horizon UHPLC system coupled to a Thermo Exploris 240 Orbitrap mass spectrometer. For LC separation, a Vanquish binary pump system (Thermo Fisher Scientific) was used with a Waters Acquity CSH C18 column (100 mm × 2.1 mm, 1.7 μm particle size) held at 35 °C under 300 μL/min flow rate. Mobile phase A consisted of 5 mM ammonium acetate in acetonitrile (ACN):H_2_O (70:30, v/v) with 125 μL/L acetic acid. Mobile phase B consisted of 5 mM ammonium acetate in isopropanol:ACN (90:10, v/v) with the same additive. For each sample run, mobile phase B was initially held at 2% for 2 min and then increased to 30% over 3 min. Mobile phase B was further increased to 50% over 1 min and 85% over 14 min and then raised to 99% over 1 min and held for 4 min. The column was re-equilibrated for 5 min at 2% B before the next injection. Samples (5µl) were injected by a Vanquish Split Sampler HT autosampler (Thermo Fisher Scientific), while the autosampler temperature was kept at 4 °C. The samples were ionized by a heated ESI source kept at a vaporizer temperature of 350 °C. Sheath gas was set to 50 units, auxiliary gas to 8 units, sweep gas to 1 unit, and the spray voltage was set to 3500 V for positive mode and 2500 V for negative mode. The inlet ion transfer tube temperature was kept at 325 °C with 70% RF lens. For targeted analysis, the MS was operated separately in positive and negative mode, with targeted scans to oxidized CoQ_10_ H^+^ adduct (m/z 863.6912), oxidized CoQ_10_ NH_4_^+^ adduct (m/z 880.7177), reduced CoQ_10_H_2_ H^+^ adduct (m/z 865.7068), reduced CoQ_10_H_2_ NH_4_^+^ adduct (m/z 882.7334), oxidized CoQ_8_ H^+^ adduct (m/z 727.566), oxidized CoQ_8_ NH_4_^+^ adduct (m/z 744.5935) and CoQ intermediate PPHB_10_ H^-^ adduct (m/z 817.6504). MS acquisition parameters include resolution of 45,000, HCD collision energy (45% for positive mode and 60% for negative mode), and 3s dynamic exclusion. Automatic gain control targets were set to standard mode. The resulting CoQ intermediate data were processed using TraceFinder 5.1 (Thermo Fisher Scientific). Raw intensity values were normalized to the CoQ_8_ internal standard and protein content as determined by BCA.

### LC-MS/MS Proteomics

Cell culture pellets were resuspended in 2% SDS containing cOmplete Protease Inhibitor Cocktail (Sigma 11697498001) and heated at 95°C for 5 min. Nucleic acids were sheared with benzonase (Sigma E1014) and samples were incubated on ice for 15 min. Protein content was quantified using a BCA assay (Pierce 23225) and 100 μg of protein were alkylated and reduced in digestion solution (10 mM TCEP, 40 mM CAA, 100 mM Tris pH 8.0) for 30 min at room temperature. Protein was subjected to single-pot, solid-phase-enhanced sample preparation (SP3) to remove detergent by incubating with magnetic carboxylated SpeedBeads (Sigma GE65152105050250). After incubation (1 hour) facilitating protein binding, beads were washed with 80% ethanol and allowed to dry. The beads were resuspended in 100 mM Tris pH 8.0 and trypsin (Promega) was added to each sample in an estimated 50:1 protein:enzyme ratio for overnight digest at 37°C. The following day, the supernatant containing tryptic peptides was collected and acidified with TFA to a pH of 2.0. The peptides were desalted by solid-phase extraction cartridges (Phenomenex) and dried under vacuum (Thermo Scientific).

Samples were resuspended in 0.2% formic acid and subjected to LC-MS analysis. LC separation was performed using the Thermo Ultimate 3000 RSLCnano system. A 15 cm EASY-Spray PepMap RSLC C18 column (150 mm x 75 mm, 3 mm) was used at 300 nL/min flow rate with a 120 min gradient using mobile phase A consisting of 0.1% formic acid in H_2_O, and mobile phase B consisting of 0.1% formic acid in ACN:H_2_O (80:20, v/v). EASY-Spray source was used and the temperature was at 35°C. Each sample run was held at 4% B for 5 min and increased to 30% B over 100 min, followed by 5 min at 99% B and back to 4% B for equilibration for 10 min. An Acclaim PepMap C18 HPLC trap column (20 mm x 75 mm, 3 mm) was used for sample loading.

MS detection was performed with a Thermo Exploris 240 Orbitrap mass spectrometer in positive mode. The source voltage was 1.8 kV, ion transfer tube temperature was set to 275°C, and RF lens was at 70%. Full MS spectra were acquired from m/z 350 to 1400 at the Orbitrap resolution of 60000, with the normalized AGC target of 300% (3E6). Data-dependent acquisition (DDA) was performed with a 3 second duty cycle with the charge state of 2-6, an isolation window width of 2, and intensity threshold of 5E3. Dynamic exclusion was 20 s with the exclusion of isotopes. Other settings for DDA include Orbitrap resolution of 15000 and HCD collision energy of 30%.

Raw files were analyzed by SequestHT Search Engine incorporated in Proteome Discoverer v.2.5.0.400 software against human databases downloaded from Uniprot. Label-free quantification was enabled in the searches.

### Blue native electrophoresis and in gel activity stains

Sample preparation and blue native electrophoresis (BNE) of cultured cell pellets were essentially done as previously described ^61,62^. Briefly: For BNE cells were collected by scraping and centrifugation, further disrupted using a pre-cooled motor-driven glass/Teflon Potter-Elvehjem homogenizer at 2000 rpm and 40 strokes. Homogenates were centrifuged for 15 min at 600 g to remove nuclei, cell debris, and intact cells. Mitochondrial membranes were sedimented by centrifugation of the supernatant for 15 min at 22,000g. Mitochondria enriched membranes from 20 mg cells were resuspended in 35 µl solubilisation buffer (50 mM imidazole pH 7, 50 mM NaCl, 1 mM EDTA, 2 mM aminocaproic acid) and solubilized with 10 µl 20% digitonin (Serva). Samples were supplemented with 2.5 µl 5% Coomassie G250 in 500 mM aminocaproic acid and 5 µl 0.1% Ponceau S in 50% glycerol. Equal protein amounts of samples were loaded on to a 3 to 18% polyacrylamide gradient gel (dimension 14x14 cm). After native electrophoresis in a cold chamber, blue-native gels were fixed in 50% (v/v) methanol, 10% (v/v) acetic acid, 10 mM ammonium acetate for 30 min and stained with Coomassie (0.025% Serva Blue G, 10% (v/v) acetic acid). BNE and immunoblotting analysis of n-dodecyl β-D-maltoside (DDM) solubilised mitochondria isolated from U2OS cells was performed as previously described ^62^. Briefly, mitochondrial membranes were solubilised using 2 mg/mg protein DDM on ice for 20 min and samples 100µg were separated on a 4 to 16% native polyacrylamide gel following manufacturers’ instructions (Novex® NativePAGE™ Bis-Tris Gel System and reagents). Separated OxPhos complexes were transferred to a PVDF membrane and immunoblotted for CI, CII and CV. In gel activity stains were performed as described in ^63^.

### Complexome Profiling

Enriched mitochondrial proteins were extracted from U2OS cells and patient fibroblasts. Mitochondrial membranes were solubilized with digitonin as described in ^64^. Equal amounts of solubilised mitochondrial protein extracts were subjected to a 3 to 18% polyacrylamide gradient gel (14cm ×14cm) for BN-PAGE as outlined in (^61^). The gel was then stained with Coomassie blue and cut into equal fractions then transferred to 96-well filter plates. The gel fractions were then destained in 50 mM ammonium bicarbonate (ABC) followed by protein reduction using 10 mM DTT and alkylation in 20 mM iodoacetamide. Protein digestion was carried out in digestion solution (5 ng trypsin/μl in 50 mM ABC, 10% ACN, 0.01% (m/v) ProteaseMAX surfactant (Promega), 1 mM CaCl_2_) at 37°C for at least 12 hours. The peptides were dried in a SpeedVac (ThermoFisher) and eluted in 1% ACN and 0.5% formic acid into a new 96-well plate. Nano liquid chromatography and mass spectrometry (nanoLC/MS) was carried out on Thermo Scientific™ Q Exactive Plus equipped with ultra-high performance liquid chromatography unit Dionex Ultimate 3000 (ThermoFisher) and Nanospray Flex Ion-Source (ThermoFisher). The data was analysed using MaxQuant software at default settings and the recorded intensity-based absolute quantifications (iBAQ) values were normalised to control cell membranes. The mass spectrometry complexomics data together with a detailed description have been deposited to the ProteomeXchange Consortium via the PRIDE partner repository ^54^ with the dataset identifier PXD055511.

## Acknowledgements

This work was supported by the Wellcome Centre for Mitochondrial Research (203105/Z/16/Z) (RNL, ZMAC-L and RWT); the Lily Foundation (RWT); Biochemical Society Vacation Scholarship funding (MO); a Barbour Foundation PhD studentship awarded to JJC; grants from the Deutsche Forschungsgemeinschaft (DFG), FOR5046 grant number WI 3728/1-1 (project number 426950122), and WI-3728/3-1 (project number 515944830) (IW), the German Federal Ministry of Education and Research (BMBF, Bonn, Germany) grant to the German Network for Mitochondrial Disorders (mitoNET, 01GM1906D) (IW) and NIH award R35GM131795 and funds from the BJC Investigator Program (DJP). MO and RWT receive additional financial support from the Pathology Society and Mito Foundation. MO is supported by Fight for Sight. RWT is also supported by the Medical Research Council (MRC) International Centre for Genomic Medicine in Neuromuscular Diseases (MR/S005021/1), the UK NIHR Biomedical Research Centre in Age and Age-Related Diseases award to the Newcastle upon Tyne Hospitals NHS Foundation, the Lily Foundation, LifeArc and the UK NHS Highly Specialised Service for Rare Mitochondrial Disorders. We thank Jana Meisterknecht for excellent technical assistance.

## Author Contributions

Conceptualization: MO, IW, DJP and RWT

Data collection, curation and formal analysis: MO, RMG, JJC, JH, KT, CRW, PCT, AC-O

Supervision: RNL, ZMAC-L, IW, DJP and RWT

Drafting the manuscript: MO, RMG, JJC, IW, DJP and RWT

Critical revision and editing of the manuscript: all authors

Funding acquisition: MO, IW, DJP and RWT

