## Supplemental Information for "RTN4IP1 is essential for the final stages of mitochondrial complex I assembly and coenzyme Q biosynthesis"

#### Supplementary Figures

##### Figure SI 1

(**A**) Western blotting analysis validating the stable expression of RTN4IP1-FLAG in the U2OS WT and $RTN4IP1^{KO}$ cell lines. WT and $RTN4IP1^{KO}$ cells were infected with a FLAG-tagged RTN4IP1-encoding lentivirus, resulting in a generation of two clones of WT and $RTN4IP1^{KO}$ cells expressing RTN4IP1-FLAG ($WT^{FLAG}{}_{C1}$, $WT^{FLAG}{}_{C2}$ and $RTN4IP1^{FLAG}{}_{C1}$, $RTN4IP1^{FLAG}{}_{C2}$). GAPDH and SDHA were used as loading controls.
(**B**) Targeted lipidomic analysis on U2OS WT and $RTN4IP1^{KO}$ cells stably expressing the indicated constructs, showing a partial restoration of total $CoQ_{10}$ and $PPHB_{10}$ levels in the RTN4IP1-deficient cells expressing RTN4IP1-FLAG. *C1* and *C2* denote two separate clones. Data shown as mean ± SD (n=3). $^{**}p<0.01$, $^{***}p<0.001$, two-sided Student's t-test.

##### Figure SI 2

**(A)** Gene Ontology of Biological Process (no redundant) showing enrichment of NADH dehydrogenase complex assembly based on proteomics analysis of $RTN4IP1^{KO}$ *versus* WT, $\log_2$ FC < -1, p value<0.05.
(**B-C**) Overexpression of RTN4IP1 can partially rescue the CI defect in *RTN4IP1*-deficient cells, as shown in (**B**) on the left panel $RTN4IP1^{KO}$*(+RTN4IP1-FLAG)* compared to WT*(+GFP),* and right panel $RTN4IP1^{KO}$*(+RTN4IP1-FLAG)*/$RTN4IP1^{KO}$*(+GFP*). In (**C**) detailed analysis of CI proteins corresponding to individual colour-coded modules is shown on the volcano plot, where empty circles represent CI subunits in $RTN4IP1^{KO}$*(+GFP*)/WT(+*GFP*) and full circles correspond to $RTN4IP1^{KO}$(+RTN4IP1-FLAG)/$RTN4IP1^{KO}$(+*GFP*). Data shown as mean (n=3), two-sided Student's t-test.

##### Figure SI 3

OxPhos complexes of *RTN4IP1* patient and control fibroblasts analysed by complexome profiling. Complexome profiling data were presented as heat map, corresponding to subunits of individual OxPhos complexes I-V. Mitochondrial complexes were solubilized with digitonin, separated on blue native gels (BNE), cut into fractions and analysed by qualitative mass spectrometry. Assignment of complexes: complex II (II); complex III dimer ($III_2$); complex IV (IV); complex V (V); small supercomplex $S_0$ of $CIII_2$ and CIV ($S_s$); supercomplex containing CI, $III_2$ and 1 copy of CIV ($S_1$) and higher order supercomlexes ($S_n$). The relative abundance of each protein was represented from low to high according to the colour scale illustrated on the bottom right.

**Figure SI 4**

CI assembly in *RTN4IP1*-derived patient fibroblasts corresponding to control cells. Complexome profiling data of CI and identified assembly factors were sorted to their assembly modules and presented as heatmap. Assignment of complexes: higher order supercomlexes ($S_n$); supercomplex $S_1$ of CI, $CIII_2$ and CIV ($S_1$); intermediate of CI including modules Q, ND1m, ND2m, ND4m and assembly factors ($I_{int}$); intermediate of Q-module containing assembly factors ($Q_{int}$) and intermediate of ND4-module containing assembly factors ($ND4_{int}$).

#### Supplementary Tables

##### Supplementary Table 1

Plasmids and oligonucleotides used in this study:

| **Plasmid** | **Description** | **Use** |
|---|---|---|
| pLVX-AcGFP1-N1 | GFP in pLVX | Cell line generation |
| pLVX-AcGFP1-N1-RTN4IP1-FLAG | RTN4IP1-Flag in pLVX | Cell line generation |

| **Primer** | **Sequence (5' to 3')** | **Use** |
|---|---|---|
| pLVX F | AAGACGATGACGACAAGTAAtgagcggccgcgactctaga | Gibson assembly |
| pLVX R | CAAGTCTTCAGAAATTCCATgaccggtggatcccgggccc | Gibson assembly |
| RNT4IP1-FLAG F | accgcgggcccgggatccaccggtcATGGAATTTCTGAAGACTTG | Gibson assembly |
| RTN4IP1-FLAG R | aattatctagagtcgcggccgctcaTTACTTGTCGTCATCGTCTT | Gibson assembly |

# SI1

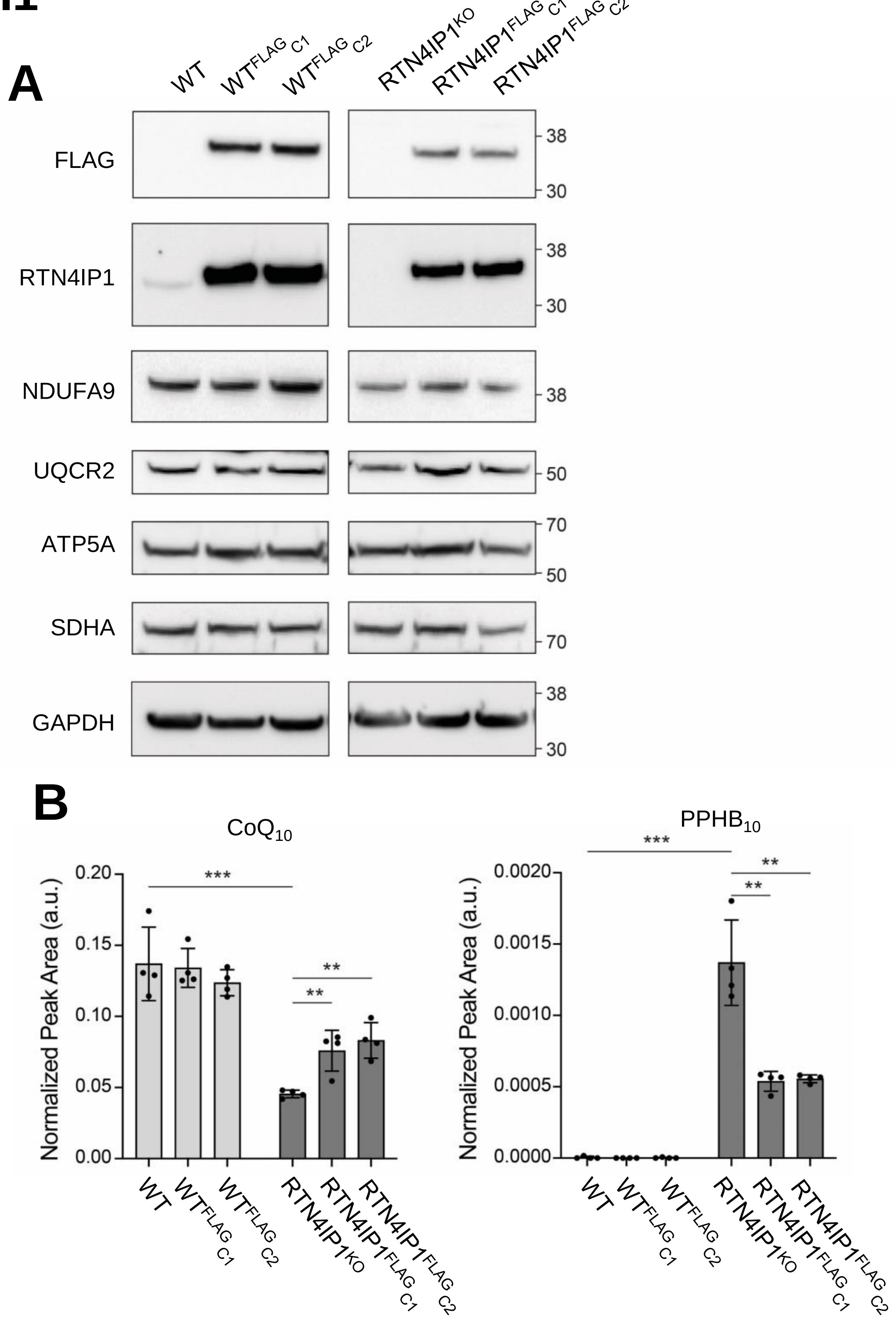
A
WT
WT FLAG C1
WT FLAG C2
RTN4IP1 KO
RTN4IP1 FLAG C1
RTN4IP1 FLAG C2
FLAG
RTN4IP1
NDUFA9
UQCR2
ATP5A
SDHA
GAPDH
38
30
50
70
B
CoQ10
PPHB10
Normalized Peak Area (a.u.)
0.20
0.15
0.10
0.05
0.00
0.0020
0.0015
0.0010
0.0005
0.0000
***
**

### Figure SI2

**A**

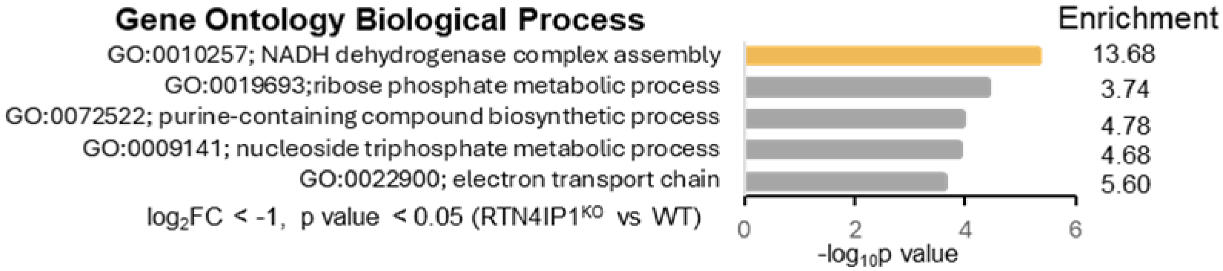


**B**

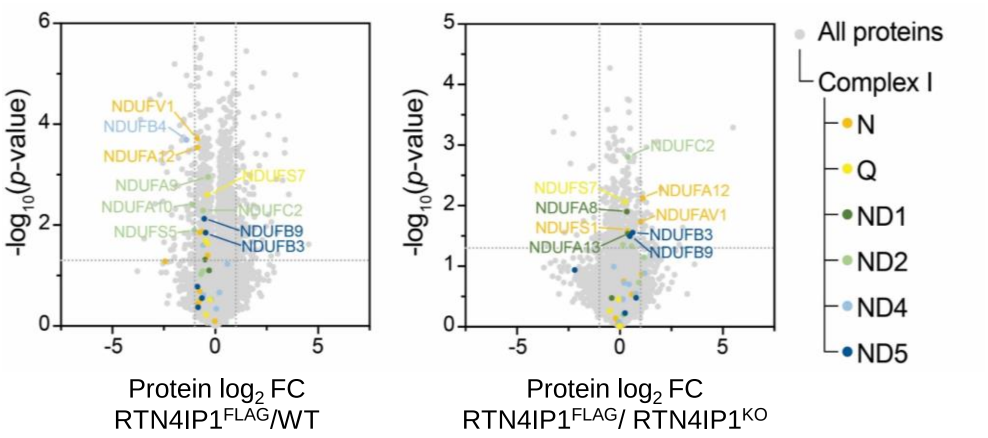


**C**

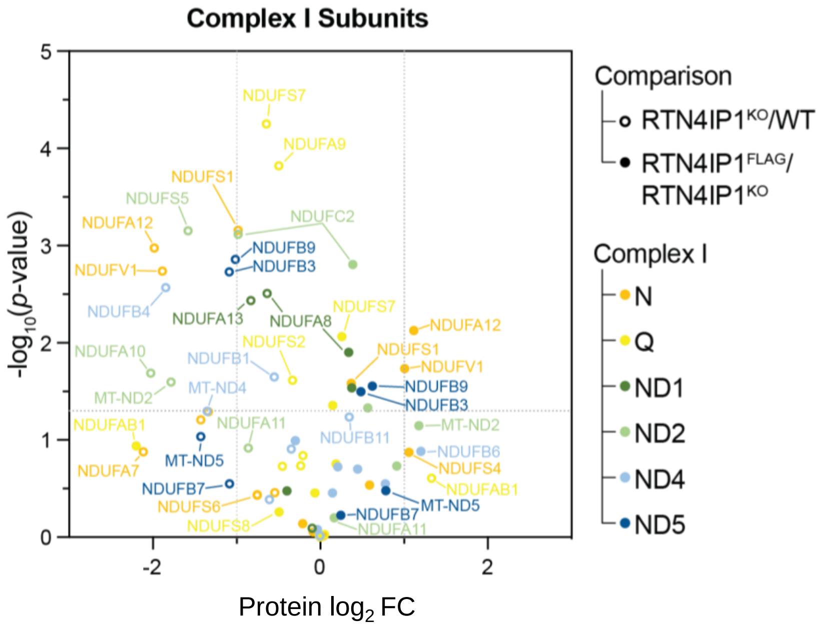

**SI3**

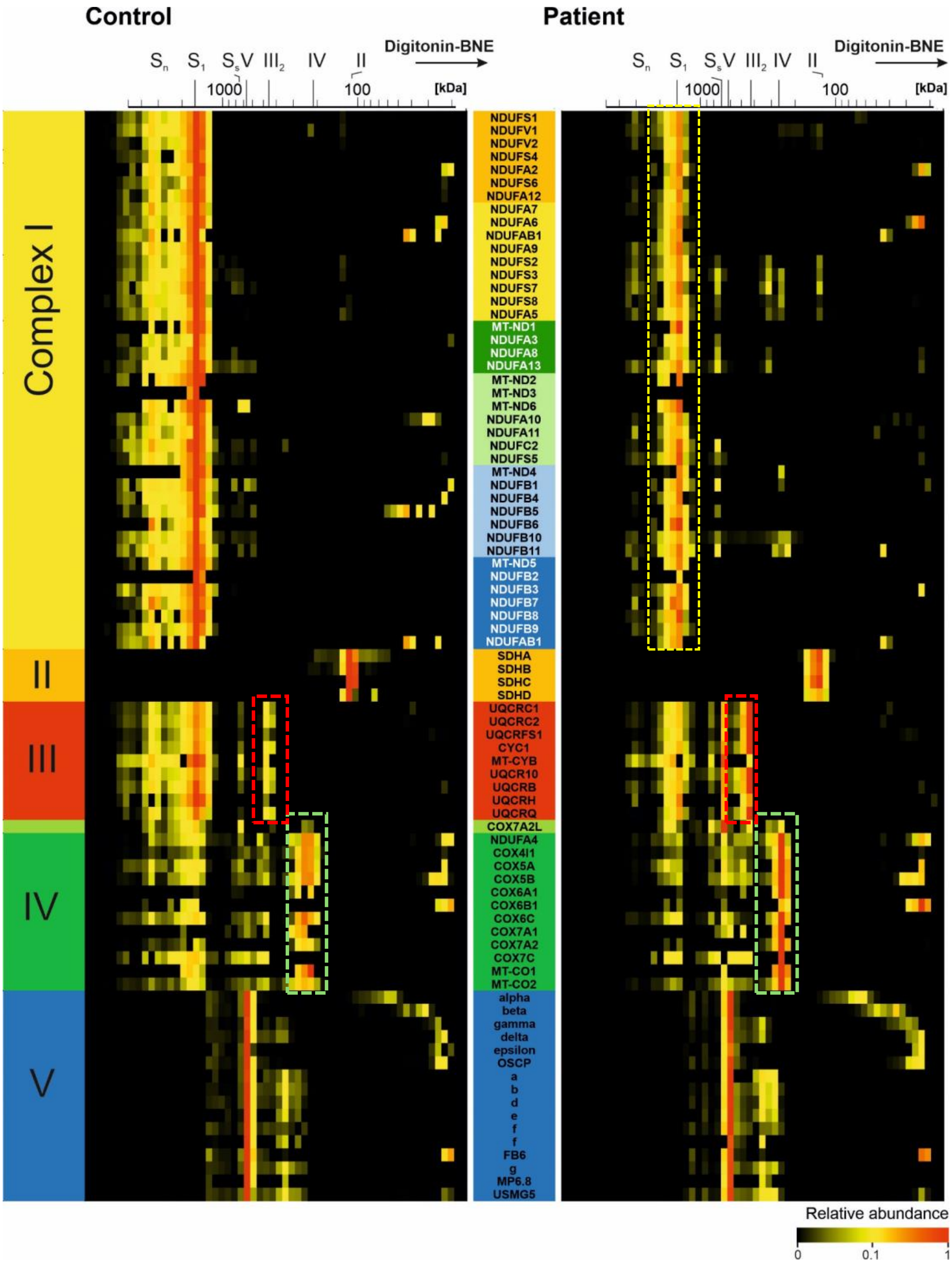

Control
Patient
Digitonin-BNE
Sn S1 SsV III2 IV II
1000
100
[kDa]
Complex I
II
III
IV
V
NDUFS1
NDUFV1
NDUFV2
NDUFS4
NDUFA2
NDUFS6
NDUFA12
NDUFA7
NDUFA6
NDUFAB1
NDUFA9
NDUFS2
NDUFS3
NDUFS7
NDUFS8
NDUFA5
MT-ND1
NDUFA3
NDUFA8
NDUFA13
MT-ND2
MT-ND3
MT-ND6
NDUFA10
NDUFA11
NDUFC2
NDUFS5
MT-ND4
NDUFB1
NDUFB4
NDUFB5
NDUFB6
NDUFB10
NDUFB11
MT-ND5
NDUFB2
NDUFB3
NDUFB7
NDUFB8
NDUFB9
NDUFAB1
SDHA
SDHB
SDHC
SDHD
UQCRC1
UQCRC2
UQCRFS1
CYC1
MT-CYB
UQCR10
UQCRB
UQCRH
UQCRQ
COX7A2L
NDUFA4
COX4I1
COX5A
COX5B
COX6A1
COX6B1
COX6C
COX7A1
COX7A2
COX7C
MT-CO1
MT-CO2
alpha
beta
gamma
delta
epsilon
OSCP
a
b
d
e
f
f
FB6
g
MP6.8
USMG5
Relative abundance
0
0.1
1

# SI4

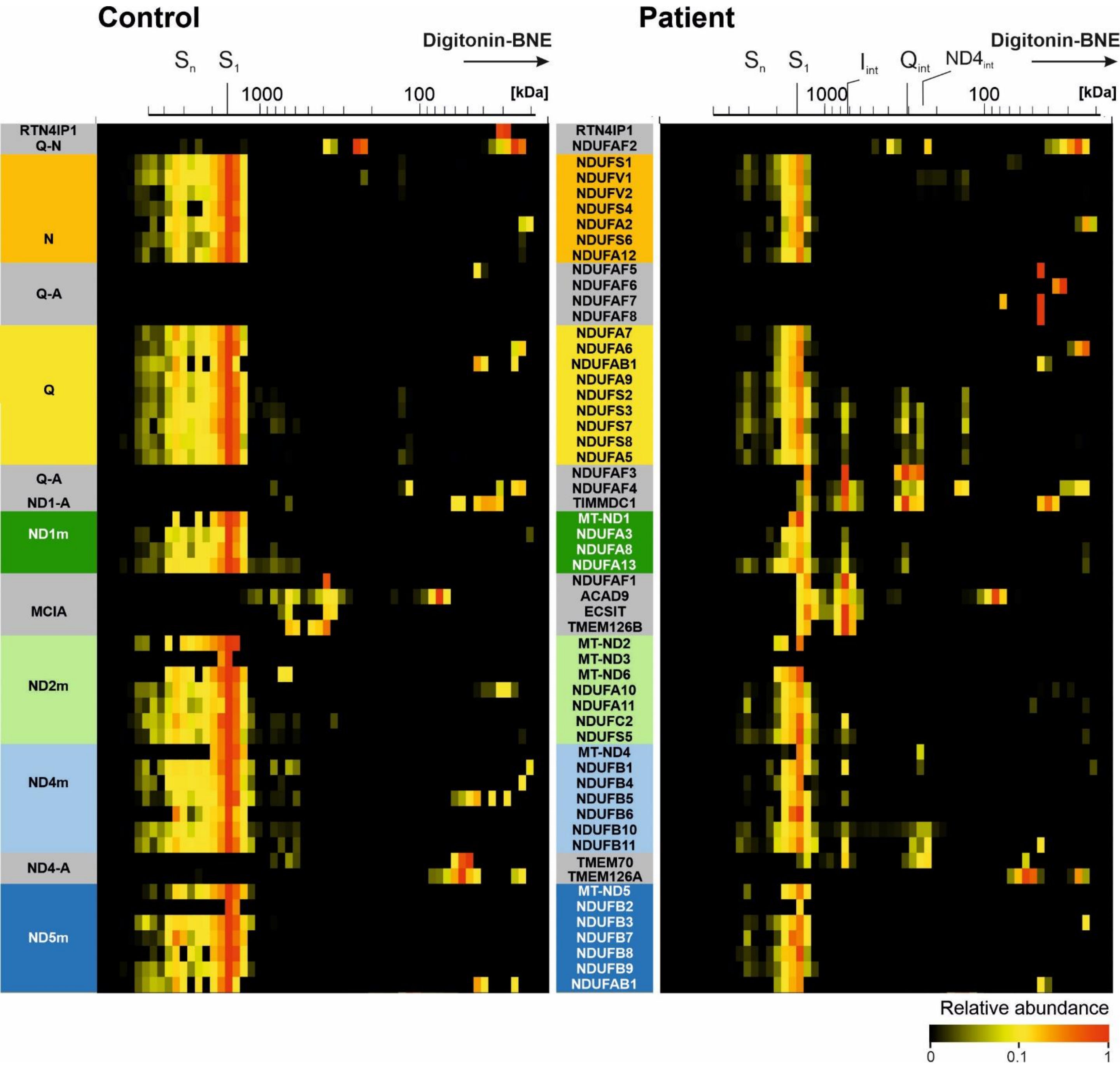
Control
Patient
Digitonin-BNE
Sn S1
Iint Qint ND4int
1000
100
[kDa]
RTN4IP1
Q-N
N
Q-A
Q
ND1-A
ND1m
MCIA
ND2m
ND4m
ND4-A
ND5m
RTN4IP1
NDUFAF2
NDUFS1
NDUFV1
NDUFV2
NDUFS4
NDUFA2
NDUFS6
NDUFA12
NDUFAF5
NDUFAF6
NDUFAF7
NDUFAF8
NDUFA7
NDUFA6
NDUFAB1
NDUFA9
NDUFS2
NDUFS3
NDUFS7
NDUFS8
NDUFA5
NDUFAF3
NDUFAF4
TIMMDC1
MT-ND1
NDUFA3
NDUFA8
NDUFA13
NDUFAF1
ACAD9
ECSIT
TMEM126B
MT-ND2
MT-ND3
MT-ND6
NDUFA10
NDUFA11
NDUFC2
NDUFS5
MT-ND4
NDUFB1
NDUFB4
NDUFB5
NDUFB6
NDUFB10
NDUFB11
TMEM70
TMEM126A
MT-ND5
NDUFB2
NDUFB3
NDUFB7
NDUFB8
NDUFB9
NDUFAB1
Relative abundance
0
0.1
1
